# Convergence of proprioceptive and visual feedback on neurons in primary motor cortex

**DOI:** 10.1101/2021.05.01.442274

**Authors:** Kevin P. Cross, Douglas J. Cook, Stephen H. Scott

## Abstract

An important aspect of motor function is our ability to rapidly generate goal-directed corrections for disturbances to the limb or behavioural goal. Primary motor cortex (M1) is a key region involved in feedback processing, yet we know little about how different sources of feedback are processed by M1. We examined feedback-related activity in M1 to compare how different sources (visual versus proprioceptive) and types of information (limb versus goal) are represented. We found sensory feedback had a broad influence on M1 activity with ∼73% of neurons responding to at least one of the feedback sources. Information was also organized such that limb and goal feedback targeted the same neurons and evoked similar responses at the single-neuron and population levels indicating a strong convergence of feedback sources in M1.

## Introduction

Sensory feedback plays a critical role in ensuring motor actions are successfully performed, providing information about motor errors due to external disturbances and internal noise inherent in the sensory and motor systems. Feedback is also essential for generating overt corrections such as when someone bumps your arm while moving, or when the behavioural goal unexpectedly moves such as a glass tipping over when the table is bumped. While vision plays a dominant role for identifying most behavioural goals, both vision and proprioception are available for feedback about the limb. Performing most motor actions thus requires combining visual feedback of the goal with feedback of the limb from proprioception and vision.

Primary motor cortex (M1) plays an important role in generating goal-directed corrections during motor actions. M1 receives rich sensory inputs from many brain regions involved in proprioceptive and visual processing including the parietal and frontal cortices (Jones et al., 1978; Zarzecki and Strick, 1978; Crammond and Kalaska, 1989; Porter and Lemon, 1993; Buneo et al., 2002; Pesaran et al., 2006; McGuire and Sabes, 2011; Bremner and Andersen, 2012; Dea et al., 2016; Omrani et al., 2016; Gamberini et al., 2017; Piserchia et al., 2017; Kalidindi et al., 2020; Takei et al., 2021), as well as input from cerebellum (Conrad et al., 1975; Vilis et al., 1976; Strick, 1983; Guo et al., 2020; Sauerbrei et al., 2020). M1 rapidly responds to proprioceptive feedback of the limb within ∼20-40ms of an applied mechanical load (Evarts and Tanji, 1976; Wolpaw, 1980; Lemon, 1981a; Suminski et al., 2009; Pruszynski et al., 2011, 2014; Omrani et al., 2014; Heming et al., 2019) and to visual feedback of the limb and goal within ∼70ms (Georgopoulos et al., 1983; Cisek and Kalaska, 2005; Ames et al., 2014; Stavisky et al., 2017). Thus, M1 receives both visual and proprioceptive feedback, but we know little about how these different sources of sensory information are organized in M1 during motor actions.

On one extreme, all three feedback sources could target a similar population of neurons (convergence hypothesis). This hypothesis is consistent with the assumption that the motor system computes a difference vector between the visual location of the goal and an estimate of hand position, which is then used to calculate motor commands (Bullock et al., 1998; Sober and Sabes, 2003; Shadmehr and Wise, 2005; Burns and Blohm, 2010). This difference vector is commonly assumed to be computed upstream in premotor and/or posterior parietal cortices (Buneo et al., 2002; Pesaran et al., 2006; McGuire and Sabes, 2011; Bremner and Andersen, 2012; Piserchia et al., 2017). Consistent with this hypothesis are studies showing how corrective responses for sensory feedback of the limb can depend on properties of the goal including its location (Brenner and Smeets, 2003; Mutha et al., 2008; Pruszynski et al., 2008; Yang et al., 2011; Dimitriou et al., 2013; Cluff and Scott, 2015). Visual feedback about the limb can also affect how participants correct for proprioceptive errors (Wei and Körding, 2008; Ito and Gomi, 2020). If a difference vector is computed upstream and transmitted to M1 during movement, the prediction is that a common group of neurons in M1 should rapidly respond to mechanical and visual disturbances of the limb as well as visual disturbances of the goal.

Alternatively, each feedback source may influence M1 independently (independence hypothesis). The motor system rapidly responds to proprioceptive (∼20-60ms) and visual (90-120ms) feedback, which may not allow the brain sufficient time to perform the necessary computations needed to integrate feedback sources. Behavioural studies suggest that the motor system may have independent representations of the limb and goal (Brenner and Smeets, 2003; Franklin et al., 2016) as well as independent representations for visual and proprioceptive feedback of the limb (Krakauer et al., 1999; Shadmehr and Krakauer, 2008; Oostwoud Wijdenes and Medendorp, 2017). M1 receives inputs from many brain areas including primary somatosensory cortex (S1; Jones et al., 1978; Dea et al., 2016), an area that is primarily involved with processing proprioceptive and cutaneous feedback. The prediction for this hypothesis is that each feedback source will influence an independent set of neurons in M1.

Here, we explored these two hypotheses by training monkeys to make goal-directed reaches while disturbances to the limb and goal were applied. Our results demonstrate that proprioceptive feedback of the limb and visual feedback of the limb and the goal influence similar groups of neurons in M1. As well, M1 activity patterns generated by each feedback source were quite similar at the single-neuron and population levels. Collectively, our results demonstrate visual and proprioceptive feedback are highly organized in M1, consistent with the convergence hypothesis.

## Results

### Behaviour, neural and muscle activities are similar with and without visual feedback of hand position

We trained monkeys to reach to a goal and on random trials applied perturbations to either the goal or limb during the movement (Figure 1A). For two perturbations, they involved either a jump to the visual feedback of the goal or visual feedback of the limb (white cursor; Figure 1B, C). We also probed proprioceptive feedback of the limb by applying a mechanical load that physically displaced the limb (Figure 1D). To isolate the proprioceptive feedback response only, we transiently removed visual feedback of the hand (white cursor, removed for 200ms) at the time of the mechanical load. In order to verify this transient removal of vision had minimal impact on performance, we compared unperturbed trials where cursor feedback was provided for the entire trial (cursor-on trials) with trials where cursor feedback was transiently removed (200ms, cursor-off trials; Figure 1A). We found cursor-on and cursor-off trials had similar movement times (Figure 1E, S1A, E), but that there was an ∼33% increase for cursor-off trials in the endpoint distance (distance the reach endpoint was from the goal; Figure S1C, G). Neural activity in M1 was also highly similar between cursor-on and cursor-off trials (Figure 2A) with activity magnitudes that were strongly correlated across neurons (Figure S2, S3A-D r>0.90) and had regression slopes near unity. Only ∼5% of neurons displayed significantly different activities between the trial types (black circles; two-sample t-test, p<0.01). Muscle activity was highly similar between cursor-on and cursor off-trials (Figure 2A, bottom row) with activity magnitudes that were highly correlated across muscle samples (Figure S3E, F) and with regression slopes near unity. Only 6% of muscle samples displayed significantly different activities for cursor-off and cursor-on trials. Thus, transient removal of visual feedback of the limb had minimal impact on motor performance during reaching and the corresponding M1 and muscle activities.

**Figure 1.**
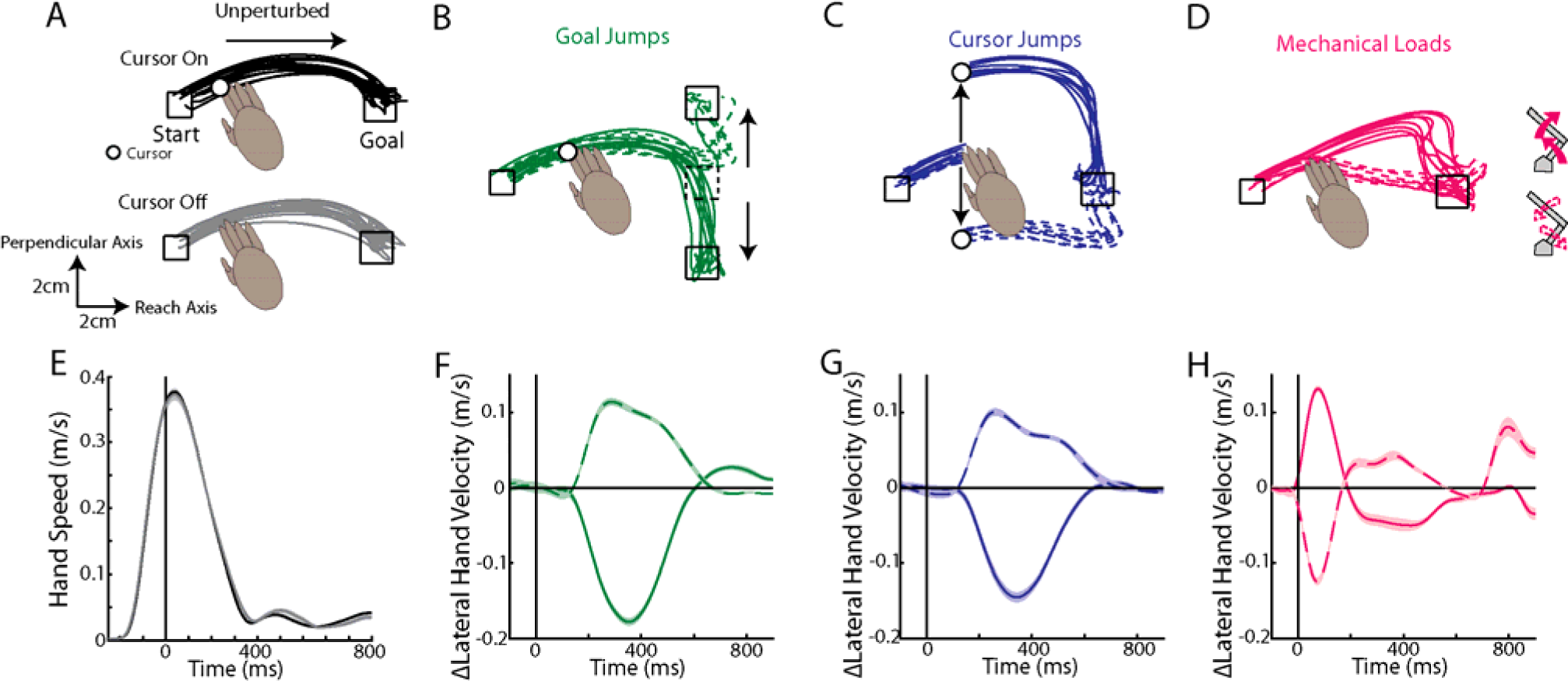
Example kinematics. A) Example hand paths of Monkey M reaching for cursor-on (top) and cursor-off trials (bottom). B-D) Example hand paths for goal jumps (B), cursor jumps (C) and mechanical loads (D). Solid and dashed lines are perturbations requiring corrections towards and away from the body, respectively. E) The average hand speed on cursor-on and cursor-off trials. F-H) The change in the lateral hand velocity for goal jumps (F), cursor jumps (G), and mechanical loads (H). Note, for the mechanical loads the change in lateral hand velocity starts at 0ms due to the displacement caused by the loads.

**Figure 2.**
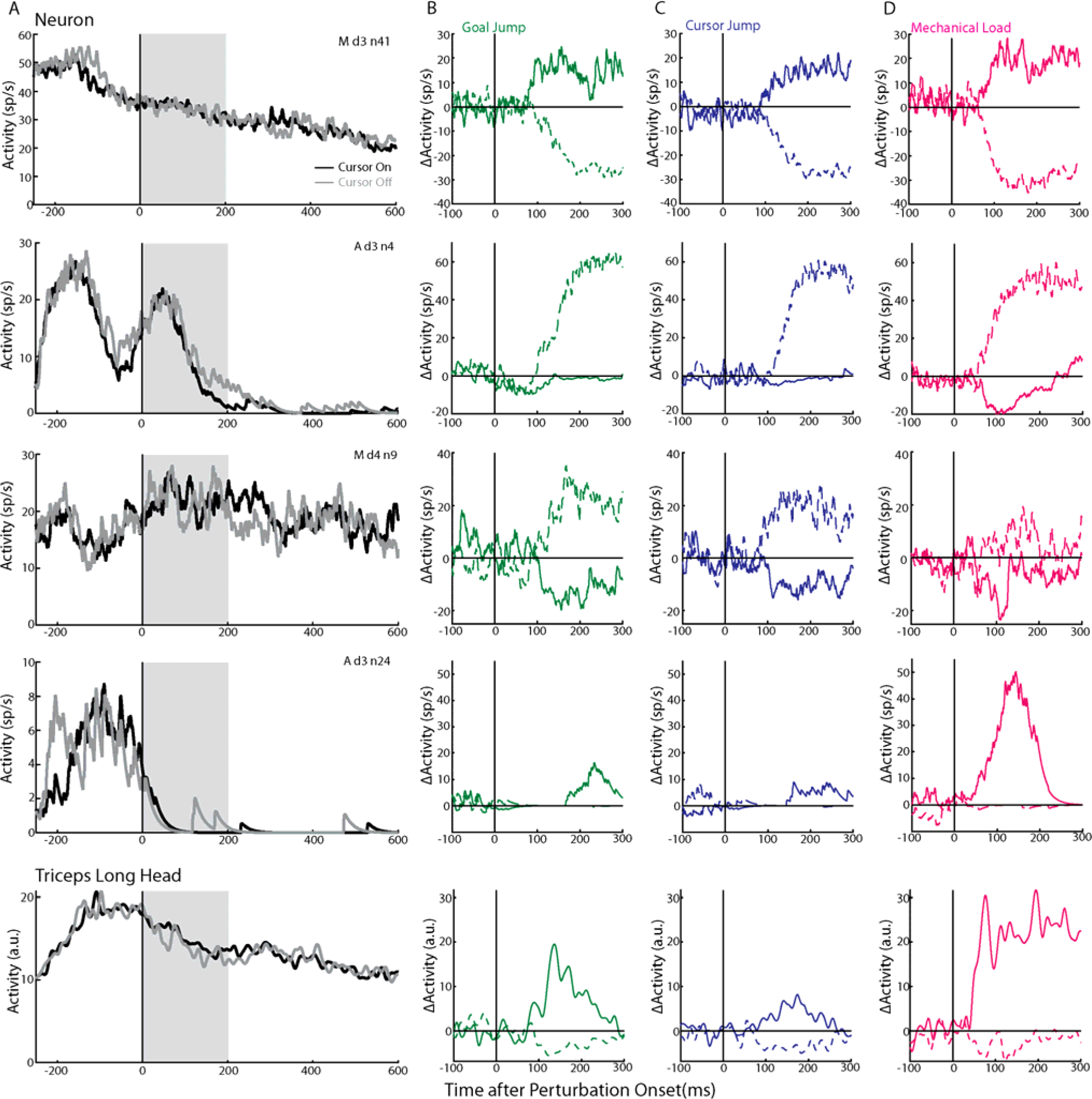
Example neuron activities. A) Activities from four example neurons (first four rows) and muscle activity (bottom row) during reaches for cursor-on (black) and cursor-off trials (grey). Grey area demarcates when vision was removed. B-D) The change in activities (ΔActivity) for the same four example neurons and muscle activity in response to the goal jumps (B), cursor jumps (C) and mechanical loads (D). Solid and dashed lines are responses to perturbations requiring corrections towards and away from the body, respectively.

### Monkeys rapidly counteract perturbations to the limb and goal

Next, we examined corrections for the different perturbation types (goal jumps, cursor jumps, and mechanical loads). Each perturbation type required corrections that moved the limb either towards the body (Figure 1B-D solid lines) or away from the body (dashed lines). Monkeys were able to quickly initiate a correction to each perturbation type within <200ms of the perturbation (Figure 1F-H). Perturbations resulted in longer movement times (24-138% increase Figure S1B, F) and greater endpoint distance (13-119% increase Figure S1D, H) than the unperturbed reaches.

Many neurons displayed robust responses following mechanical and visual perturbations with four example neurons shown in Figure 2B-D. The first neuron (Figure 2B, top row, Md3n41) displayed a reciprocal response for goal jumps within 100ms of the jump onset with an increase (solid) and decrease (dashed) in activities for corrective movements towards and away from the body, respectively. These changes in activity plateaued within 150ms of the jump onset and remained relatively constant over the next 150ms. However, the plateau for the inhibition response may reflect that the activity of the neuron was approaching 0sp/s (see Figure 2A top row). This neuron displayed a similar pattern of responses for cursor jumps (Figure 2C, top row) and mechanical loads (Figure 2D, top row). Neuron 2 (second row, Ad3n4) displayed similar excitations for corrections away from the body across the different perturbation types. Neuron 3 (third row, Md4n9) displayed a similar pattern of responses across the two visual perturbations with an increase and decrease in activities for the corrective movements away from and towards the body, respectively. This neuron had similar selectivity for the mechanical loads, however, its responses were noticeably smaller. In contrast, neuron 4 (fourth row, Ad3n24) exhibited considerably larger activity for the mechanical loads than either cursor jump or goal jump while still maintaining the same selectivity across perturbation types.

### Each perturbation type targets similar neurons in M1

Our objective is to identify whether each feedback source targeted independent groups of neurons in M1. We classified neurons that had a significant response to each perturbation type by applying a three-way ANOVA with time epoch (two levels: baseline=100ms before perturbation onset, perturbation=0-300ms after perturbation onset), perturbation type (three levels: mechanical, cursor, goal) and perturbation direction (two levels: towards and away from the body) as factors. For Monkeys M|A, we found 71|76% (n=122|65) of neurons had a significant main or interaction effect(s) with time (p<0.0125), which we labeled as perturbation-responsive neurons. We identified neurons that were responsive to a particular perturbation type by using a two-way ANOVA with time and perturbation direction as factors. Similar percentages of neurons were responsive for goal jumps (55|54%, n=94|51), cursor jumps (44|60% n=75|51) and mechanical loads (55|60% n=94|46). These neurons received sensory feedback rapidly as the onset of perturbation-related activity at the population level occurred within <100ms with responses to the mechanical loads arising earlier (Monkey M|A: 43|57ms) than for either visual jump (goal=78|74ms, cursor=83|82ms; Figure 3A, C). Similar results were found when examining individual onsets (Figure 3B, D) and a one-way ANOVA with onset type as a factor (3 levels: mechanical, goal and cursor) revealed a significant main effect (Monkey M: F(2,295)=12.6, p<0.001, Monkey A: F(2,168)=10.3, p<0.001). Post-hoc tests confirmed that onsets for the mechanical-related activity started earlier (Monkey M|A mean 119|106ms) than either visual perturbation (goal 140|143ms p=0.01|p=0.002, cursor 159|155ms p<0.001|p<0.001). Onset differences between the two visual perturbations were not significant (p=0.05|p=0.49).

**Figure 3.**
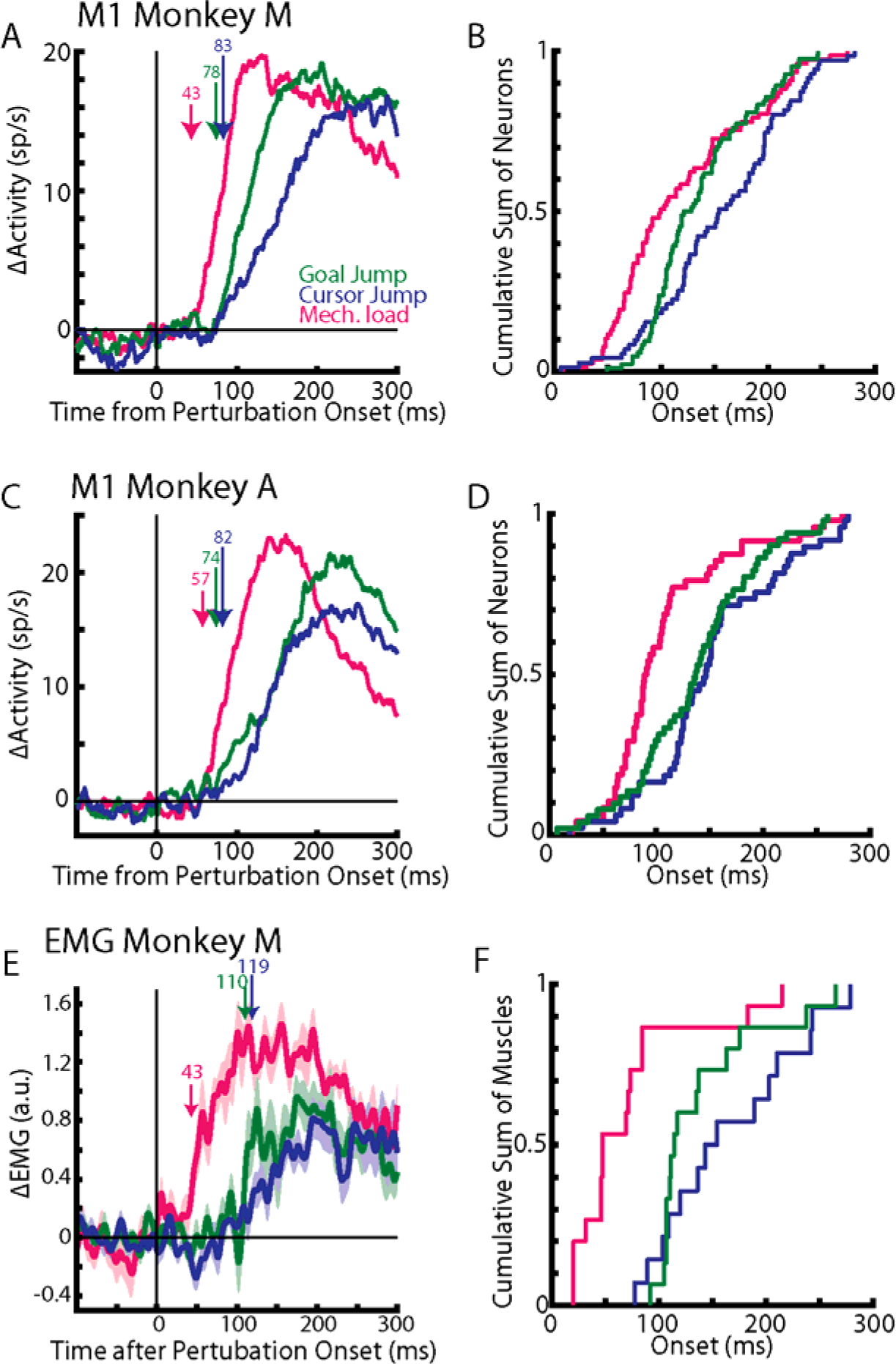
Proprioceptive feedback alters M1 activity earlier than visual feedback. A) The average activity across neurons for Monkey M. Arrows indicate when a significant increase from baseline was detected. Only neurons with significant activity for at least one perturbation type were included. B) The onset across individual neurons for each perturbation type presented as a cumulative sum. C-D) Same as A-B) except for Monkey A. E-F) Same as A-B) except for muscle activity from Monkey M.

From the percentages of neurons that responded to each perturbation type we estimated the number of neurons expected to respond to zero, one, two and three perturbation types assuming responses were independently assigned (expected distribution). Perturbation responses were significantly more overlapped than the expected distribution (Monkey M|A: χ^2^ =113.9|68.1, df=4, p<0.001|<0.001). In Monkey M|A, 15|13% (n=26|11) of neurons responded to only one perturbation type, which was 2.4|2.4 times smaller than the expected distribution (Figure 4A, C). In contrast, 28|36% (49|31) of neurons responded to all three perturbation types (common neurons), which was 2.6|3.4 times greater than the expected distribution.

**Figure 4.**
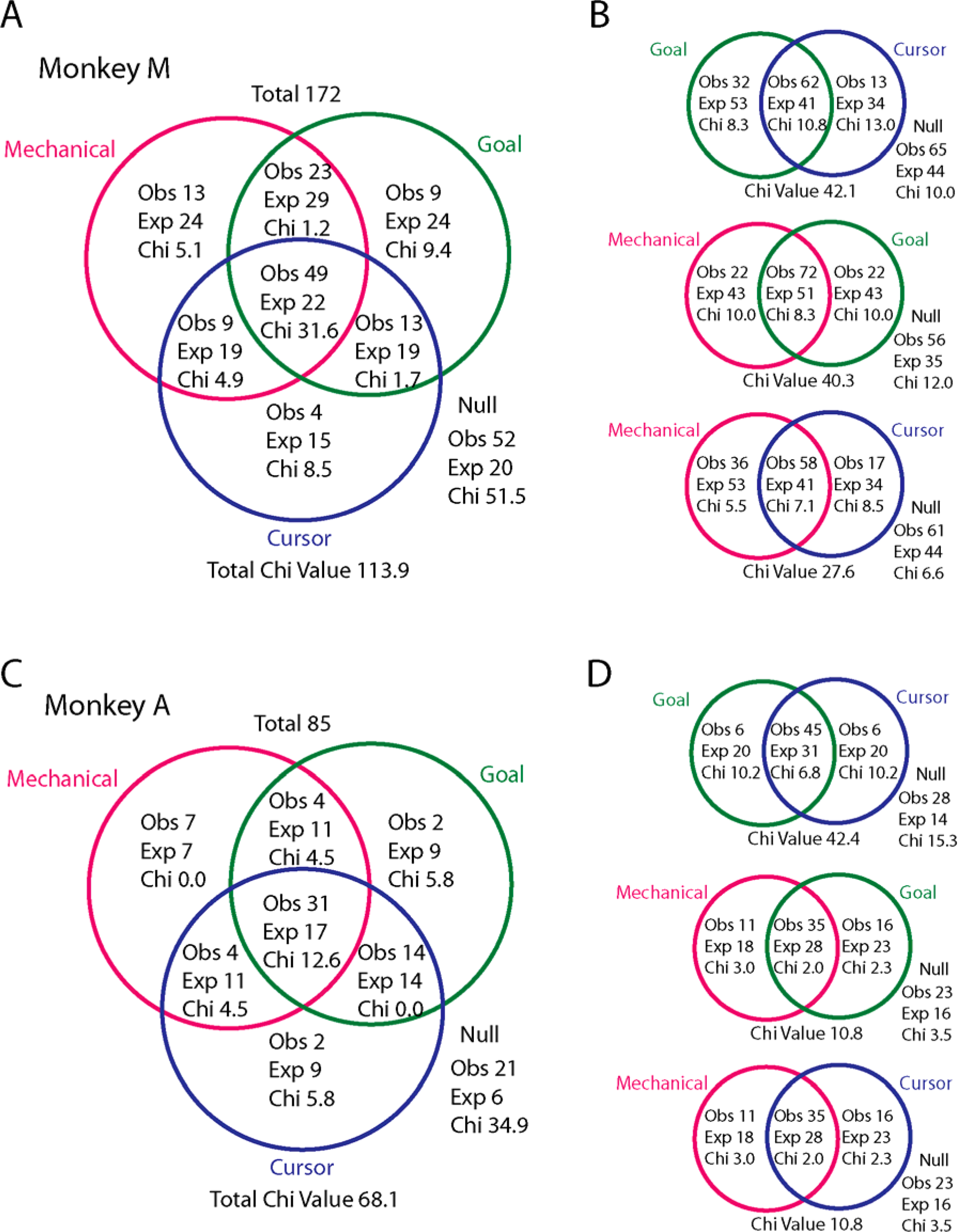
Each perturbation type influences overlapping neurons. A) Venn diagram showing the number of neurons observed (Obs) in each class for Monkey M. The diagram also shows the number of expected (Exp) neurons assuming an independent distribution. Chi reflects the classes contribution to the total χ^2^ value ([Obs-Exp]^2^/Exp). B) Venn diagrams classifying neurons using only two perturbation types for Monkey M. C-D) Same as A-B) except for Monkey A.

Thus, there was substantial overlap between groups of neurons responsive to each feedback source. However, this finding may reflect a strong overlap between just two of the perturbation types or it could reflect an overlap among all three perturbation types. We repeated the analysis across pairs of perturbation types (Figure 4B, D). Consistently, the number of neurons that responded to both perturbation types was 1.3-1.5 times greater than the expected distribution. In contrast, the number of neurons that responded to only one perturbation type was 1.5-3.4 times smaller than the expected distribution. Significant differences between the observed and expected distribution of neurons were found across all perturbation pairs (χ^2^ test, p<0.01). Collectively, these results indicate that each perturbation type influenced an overlapping set of neurons in M1.

### Neurons maintain their response ranges across perturbation types

A different way that each feedback source could independently influence M1 is by driving distinct activity patterns in the same neuron population. For example, a neuron may be strongly driven by one perturbation type but only weakly driven by a different perturbation type. At the extreme, neurons may even change their selectivity (i.e. tuning) for the loads: increase activity for the correction towards the body for one perturbation type but decrease activity for the same correction for a different perturbation type.

We explored this by examining the response range, which was calculated by taking the difference between activities for the two opposite perturbation directions (e.g. Figure 2B dashed subtracted from solid) and averaging the difference over the perturbation epoch. Neurons with greater responses for the corrections away from or towards the body will have positive or negative response ranges, respectively. Figure 5A and D compares the response ranges for goal- (abscissa) and cursor-related (ordinate) activities. Neurons responsive to all three perturbation types (black circles) resided near the unity line (solid line) and were highly correlated across the population (Monkey M|A: correlation coefficient r=0.90|0.97, p<0.001 for both). The axes that captured the largest amount of variance (dashed black lines, total least squares regression) had a slope slightly less than unity (0.84|0.86) indicating that the responses for the cursor jumps were ∼15% smaller than the goal jumps (shuffle control p=0.002|p<0.001). We found significant but noticeably weaker correlations when comparing the response ranges between the mechanical-related activities (abscissa) and activities related to either visual perturbation (ordinate; Figure 5B-C, E-F; mechanical with goal r=0.85|0.86, mechanical with cursor r=0.75|0.86, p<0.001 for all). The slope was less than unity (mechanical with goal slope=0.86|0.85, mechanical with cursor slope=0.68|0.72) indicating that the responses for the visual perturbations were ∼22% smaller than for the mechanical loads. Inclusion of all perturbation-responsive neurons yielded similar results (Figure 5 grey circles).

**Figure 5.**
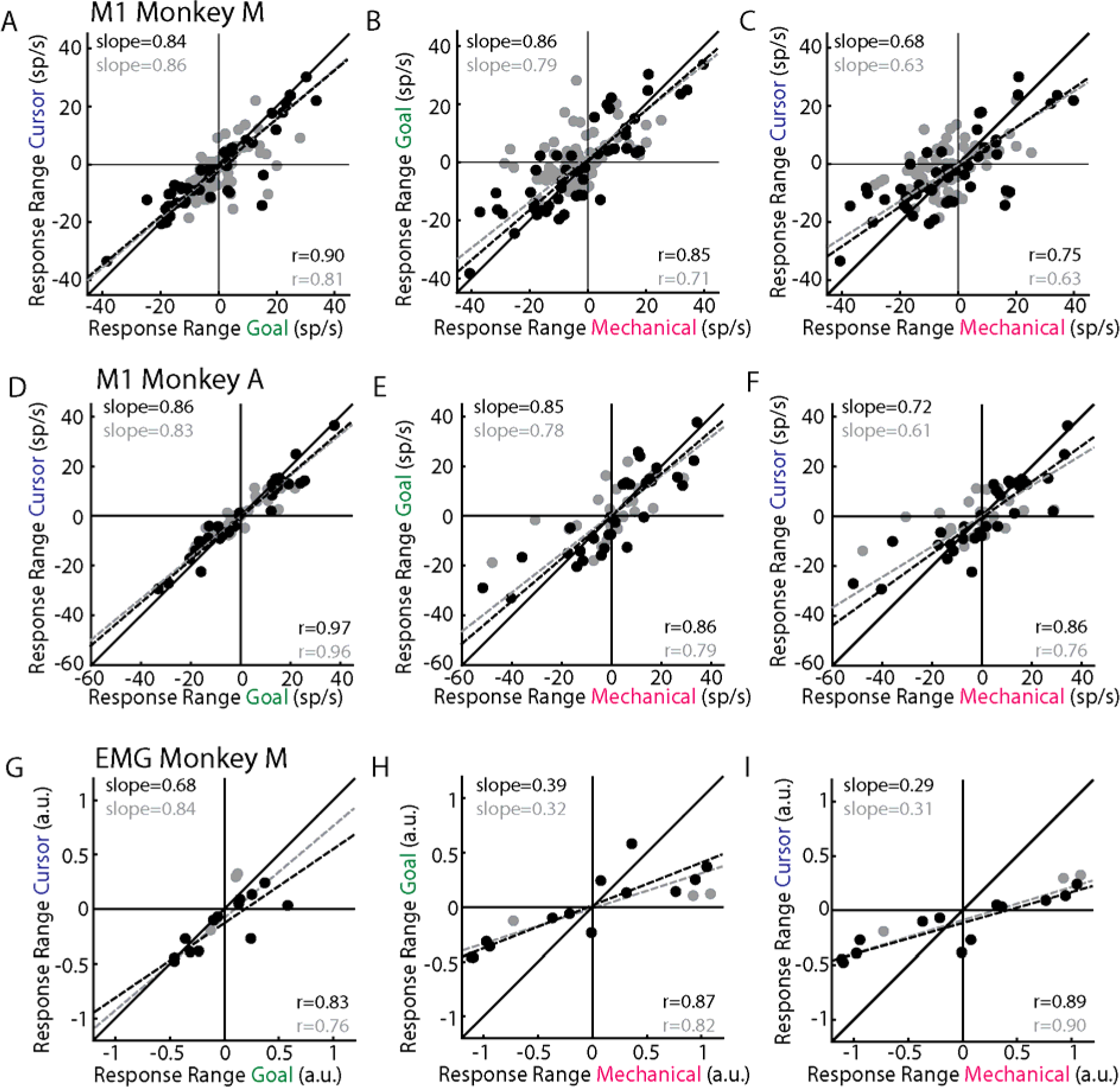
M1 neurons have similar response ranges across perturbation types. A) Comparison of the response ranges between activities for the goal and cursor jumps. Black circles: neurons responsive to all three perturbation types. Grey circles: neurons responsive to at least one perturbation type. “r” is the Pearson’s correlation coefficient. Dashed lines reflect the line of best fit identified using total least squares regression (slope indicated in quadrant 2). B) Same as A) except comparing mechanical loads and goal jumps. C) Same as A) except comparing mechanical loads and cursor jumps. D-F) Same as A-C) except for Monkey A. G-I) Same as A-C) except for muscle activity from Monkey M.

From the response ranges, we could determine if neurons maintained their selectivity for corrective movements across perturbation types. These neurons resided in the first and third quadrants of Figure 5 and we found a large majority of neurons maintained their selectivity across all three perturbation types (neurons responsive to all three perturbation types: Monkey M|A 82|87%; all perturbation-responsive neurons: 70|72%). Collectively, these results indicate that each feedback source had similar influences on individual M1 neuron responses.

Next, we compared the size of the perturbation-related activity relative to the movement-related activity during unperturbed reaching (Figure 2, S4A). Figure S4B compares the magnitude of the movement-related activity during unperturbed reaching (aligned to movement onset: movement epoch - 50 to 250ms after movement onset) with the magnitude of the response range for perturbed reaches. We found approximately equal number of neurons had either larger perturbation-related activities or movement-related activities (Figure S4B, C). Thus, the perturbation-related activity was comparable in magnitude to the activity required to generate the initial reaching movement.

### Overlap between mechanical- and visual-related M1 activity patterns at the population level

Our results so far demonstrate that each feedback source targets a largely overlapping population of M1 neurons and that individual neuron responses are generally similar across feedback sources. However, recent studies have demonstrated that the same neuron population can represent different types of information independently by sequestering the information into orthogonal subspaces (Kobak et al., 2016; Ames and Churchland, 2019; Heming et al., 2019; Keemink and Machens, 2019; Cross et al., 2020). For example, neurons in M1 have similar tuning for reach direction during preparation and execution (Crammond and Kalaska, 2000). However, these activity patterns reside in orthogonal subspaces (Kaufman et al., 2014; Elsayed et al., 2016). Thus, for the independent-input hypothesis each perturbation type may evoke an activity pattern that resides in an orthogonal subspace with respect to the other two perturbation types.

We explored this hypothesis by using principal component analysis (PCA) to identify the low-dimensional subspace each perturbation-related activity resided in. We used a cross-validated approach to prevent overestimating differences between subspaces due to sampling noise. The top-ten principal components captured 81-90% of the variance for the data used to train the principal components (open circles Figure 6A-C, E-G). Figure 6A, E shows the variance captured by the top-ten principal components generated from the goal-related activity. These components captured a substantial amount of the goal-related variance from the left-out trials (variance accounted for: Monkey M|A =55|73%) and the cursor-related variance (44|65%). These components also captured a substantial amount of the mechanical-related variance (36|43%), though noticeably smaller than either visual perturbation. Similarly, Figure 6B and F shows the variance captured by the top-ten cursor principal components. These components captured more cursor-related (49|69%) and goal-related (44|66%) variance than mechanical-related variance (30|45%). Lastly, Figure 6C, G shows the variance captured by the top-ten mechanical principal components. These components captured more mechanical-related variance (59|74%) than variance for either visual perturbation (goal 35|40%, cursor 32|40%).

**Figure 6.**
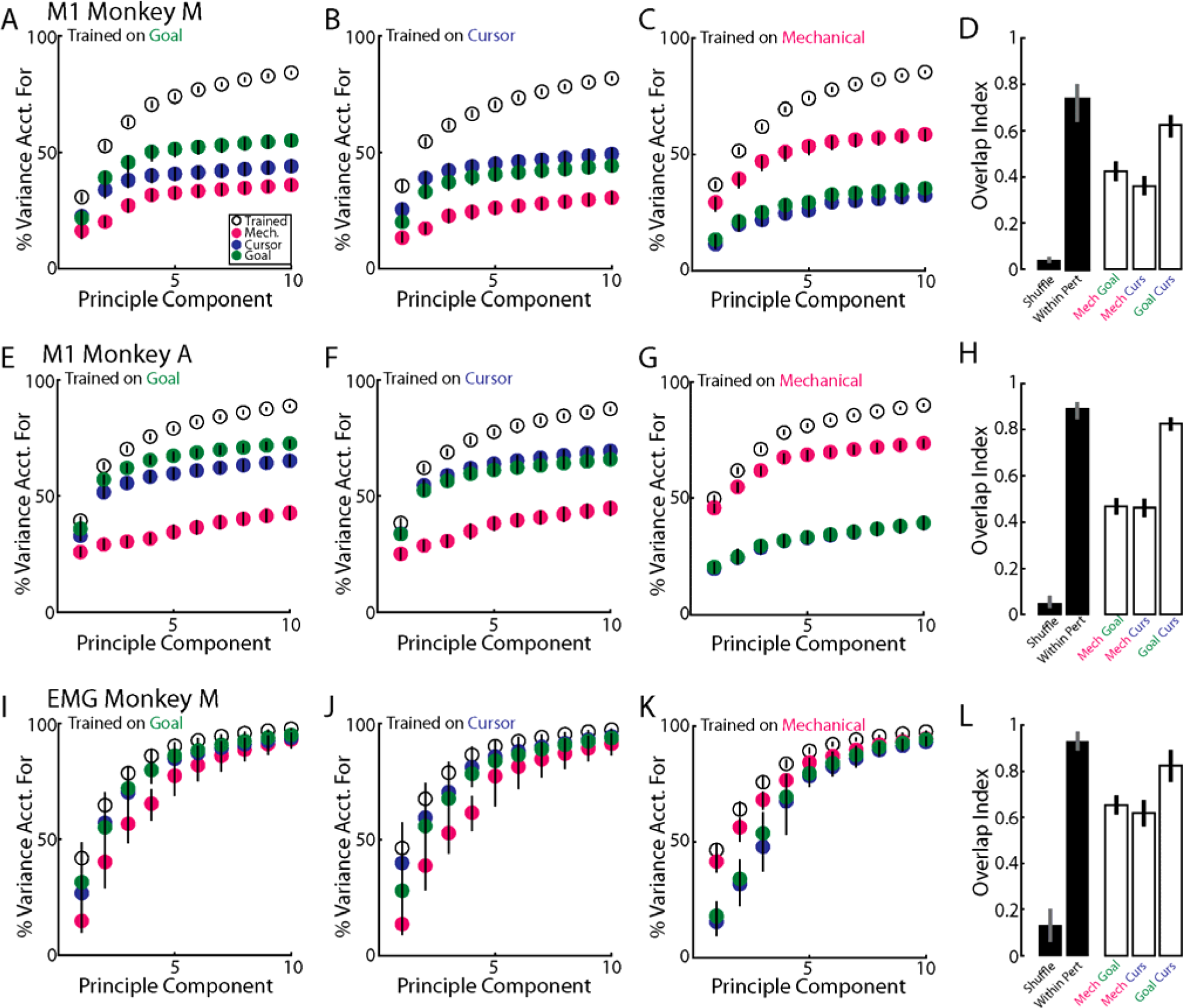
Activity patterns overlap across perturbation types. A) Variance accounted for by the top goal-jump principal components for Monkey M. Variance for the goal-jump trials was calculated for the training set (open) and for the left-out trials (red). Circles and bars denote the median and the 5^th^ and 95^th^ percentiles of the distributions. B-C) Same as A) for cursor jumps and mechanical loads. D) Overlap index between perturbation types (clear bars) and the shuffle and within-perturbation distributions (filled bars). Bars denote the median and 5^th^ and 95^th^ percentiles of the distribution. E-H) Same as A-D) except for Monkey A. I-H) Same as A-D) except for EMG from Monkey M.

Another approach to quantify the similarity in the population structure between feedback sources is by calculating the overlap index (Rouse and Schieber, 2018). The overlap index ranges from 0, indicating no overlap between subspaces (i.e. orthogonal), to 1 indicating perfect overlap. For comparison, we generated a null distribution that compared how overlapping two subspaces were after randomly shuffling neuron labels (Shuffle). We also generated a null distribution that quantified the maximum overlap expected given sampling noise by calculating the overlap between two independent samples from the same perturbation type (within-perturbation distribution). The overlap between goal- and cursor-related activities was large (Monkey M|A=0.63|0.82; Figure 6D, H) and was close to the within-perturbation distribution (0.73|0.89), though it was still significantly smaller (p=0.03|0.01). The overlap between the mechanical-related and visual-related activities were smaller than the within perturbation distribution (mechanical with goal = 0.42|0.47; mechanical with cursor = 0.36|0.46; within-perturbation p<0.001 for all), however they were still significantly greater than the shuffled distribution (p<0.001). Collectively, these results indicate each perturbation type evoked similar population-level structure.

### Overlap across perturbation types emerges rapidly with perturbation-related activity

Next, we examined how the overlap evolved over time between the different perturbation types. One possibility is that each feedback source is initially represented independently by the motor system before being gradually integrated (Franklin et al., 2016; Oostwoud Wijdenes and Medendorp, 2017). Thus, the prediction is that the overlap between perturbation types should gradually emerge. We calculated the overlap index every 20ms over the perturbation epoch (Figure S5A-F). We found the overlap index between the goal- and cursor-related M1 activities emerged within ∼100ms (Figure S5A, D, black line) post-perturbation and was comparable to the within-perturbation distributions of the goal-related (green line) and cursor-related activities (blue line). Further, the overlap between the mechanical- and visual-related M1 activities emerged within ∼100ms of the perturbation onset (Figure S5B-C, E-F). Note, that the increase in the overlap index proceeded the within-perturbation onset for the mechanical loads (red line) reflecting that M1 responds earlier for mechanical loads than visual jumps (Figure 3A, C). Interestingly, there was a small delay in the overlap between the mechanical and visual perturbations for Monkey A (Figure S5E, F) which may reflect a small-time window of integration. Similar trends were found in the muscle activity (Figure S5G-I). Thus, the overlap between perturbation types emerged rapidly in the network.

### Muscle activity exhibits similar overlap between perturbation types as M1 activity

Next, we examined the change in muscle activity in response to the different perturbation types. We found a significant change in muscle activity (Figure 2B-D bottom row) in 81% (n=13), 88% (14) and 100% (16) of muscle samples for the goal jumps, cursor jumps and mechanical loads, respectively. There was a strong correlation between response ranges for the goal- and cursor-related activities (r=0.83, p<0.001, Figure 5G) and the slope was less than unity (slope=0.68) indicating responses for the cursor jump were 32% smaller than for the goal jump. We also found strong correlations between the mechanical-related response ranges and the response ranges for either type of visual disturbance (Figure 5H-I; mechanical with goal r=0.87, mechanical with cursor r=0.89, p<0.001 for both). However, we found the slopes were considerably smaller than unity (mechanical with goal: 0.39; mechanical with cursor: 0.29) indicating that muscle activity for the visual perturbations were ∼66% smaller than for the mechanical loads. As expected, almost all (except one) of the muscle recordings maintained their selectivity across all perturbation types.

Figure 6I shows the top-ten goal principal components for muscle activity. Unlike neural activity, these ten components captured nearly all of the variance for the goal jump, cursor jump and mechanical loads. This is due to the smaller number of muscles recorded as the entire space of muscle patterns occupies a maximum of 16 dimensions. In contrast, neural activity can occupy 172 and 85 dimensions for Monkeys M and A, respectively. We mitigated this problem by restricting our observations to the top-three components as three components captured a similar amount of variance from the training data (range: 82-84%) as the ten components captured for the neural activity (82-90%). We found the top-three goal principal components captured a substantial amount of the goal- (76%) and cursor-related (74%) muscle variance but captured slightly less of the mechanical-related variance (68%). Similarly, the top-three cursor principal components captured a substantial amount of the cursor- (77% Figure 6J) and goal-related (73%) muscle variance but captured less of the mechanical-related variance (61%). Lastly, the top-three mechanical principal components captured a substantial amount of the mechanical-related muscle variance (84% Figure 6K) but captured less of the muscle variance for either visual perturbation (goal 59%, cursor 58%).

We computed the overlap index between muscle responses and found results that were similar to M1 activity (Figure 6L). There was a high overlap between the goal and cursor-related activities (0.82) that was comparable to the within-perturbation distribution (0.93), though still significantly smaller (p=0.02). We also found a partial overlap between the mechanical-related activity and the visual-related activities (mechanical and goal 0.65, mechanical and cursor 0.62), which were significantly greater than the shuffle distribution (overlap=0.13, p<0.001). Collectively, these analyses indicate that different patterns of muscle activity were needed to correct for each perturbation type which could explain the partial overlap observed between the mechanical- and visual-related M1 activities.

### Overlap is still present when examining other movement directions

One concern is whether we adequately characterized M1’s responses to each perturbation type as we sampled from only two perturbation directions. This seems unlikely as previous work has shown that a greater proportion of M1 neurons respond maximally to perturbations that involve either combined shoulder flexion and elbow extension (whole-arm extension for corrections away from body) or combined shoulder extension and elbow flexion (whole-arm flexion for corrections towards the body; Cabel et al., 2001; Scott et al., 2001; Kurtzer et al., 2006; Lillicrap and Scott, 2013). Nonetheless, we verified that sampling from more perturbation directions yielded virtually the same overlap. Monkeys completed separate blocks of the same lateral reach (Figure S6A) and also blocks of a sagittal reach starting from near the body and reaching to a distant goal (Figure S6B). For the sagittal reach, the perturbations required a corrective movement that either flexed the shoulder and elbow joints (Figure S6B solid lines) or extended the shoulder and elbow joints (dashed lines). The perturbations for the lateral and sagittal reaches yielded four perturbation directions for each perturbation type. We found response ranges were correlated between perturbation types with the strongest correlation between goal jumps and cursor jumps (Figure S6C, E, response range for sagittal reach shown only, Monkey M|A n=82|45). For the sagittal reach, activity related to goal jumps tended to be larger than activity related to cursor jumps or mechanical loads. Critically, we found the overlap between goal- and cursor-related activities was substantial (Monkey M|A=0.72|0.75, Figure S6D, F) and was close to the within-perturbation distribution (0.80|0.85), though it was still significantly smaller (p=0.01|<0.001). The overlap between the mechanical-related activity with either visual-related activity was smaller than the within-perturbation distribution (mechanical with goal = 0.50|0.49; mechanical with cursor = 0.48|0.45; within-perturbation p<0.001 for all). However, it was still significantly greater than the shuffled distribution (p<0.001).

### M1 is ∼3 times more sensitive to proprioceptive than visual feedback

So far, we have compared visual perturbations that instantaneously jump the position of the goal or cursor, with mechanical perturbations that gradually displaced the limb over 100-200ms (Figure 1H). While cursor and target jumps are standard experimental techniques to assess visual feedback (Georgopoulos et al., 1983; Dimitriou et al., 2013; Ames et al., 2014; Franklin et al., 2016; Stavisky et al., 2017), the different spatial and temporal characteristics of these perturbations make it difficult to directly compare M1’s sensitivity to proprioceptive and visual feedback errors. For a direct comparison, we compared M1’s sensitivity to the mechanical loads with cursor perturbations that slid along a pre-specified trajectory (cursor slide Figure 7A-B). The cursor’s trajectory on cursor-slide trials was highly similar to the limb’s trajectory following a mechanical load for the first 200ms with an average goodness of fit (R^2^) of 0.95 and 0.93 for Monkeys M and A, respectively (Figure 7C). We found movement times for the mechanical loads were significantly shorter than for cursor slides (Figure 7D; Mann-Whitney U test, Monkey M: U=14649, n=230, p<0.001, Monkey A: U=2454, n=98, p<0.001).

**Figure 7.**
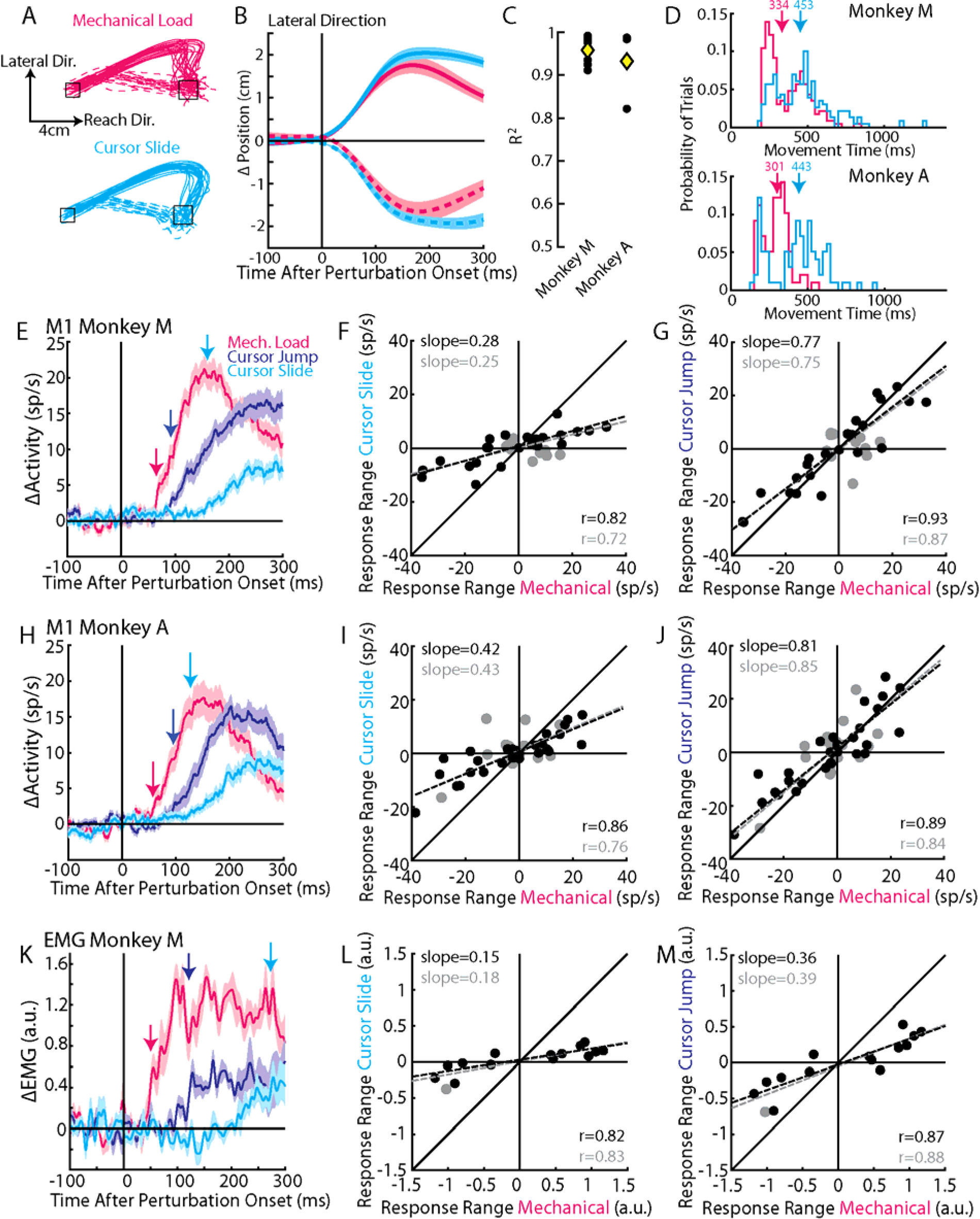
M1 is more sensitive to mechanical than visual perturbations. A) For Monkey M, hand paths for the mechanical loads (red traces) and the cursor’s path on cursor slide trials (cyan traces). B) In the lateral direction (see A), the change in position of the hand and cursor on mechanical load and cursor slide trials respectively. C) The R^2^ across sessions comparing how well the cursor slide trajectory fit the limb trajectory on the mechanical load trials (Monkey M|A n=7|3). Yellow diamonds reflect the mean. D) Movement times for all mechanical load and cursor slide trials. Arrows denote medians. E) The average activity across neurons for each perturbation type. F) Comparison of response ranges between mechanical loads and cursor slide. Presented the same as in Figure 5. G) Same as F) except for comparing mechanical loads with cursor jumps. H-J) Same as E-G) except for Monkey A. K-M) Same as E-G) except for muscle activity.

We included cursor-jump trials to identify neurons that were sensitive to visual stimuli (kinematics not shown). Note, we only used cursor perturbations to limit the number of trials as cursor and goal jumps evoked highly similar activity patterns and only differed in magnitude by ∼15% (Figure 5A, D). We recorded from 60 and 68 neurons from Monkey M and A, respectively. We found 57|57% (n=34|39) and 43|60% (26|41) responded to the mechanical loads and cursor jumps, respectively, and 40|44% (24|30) responded to both perturbations. We found the cursor slide evoked a more gradual response in M1 as compared to the mechanical load or a cursor jump (Figure 7E, H). Response ranges indicated that activity related to the cursor slide was ∼65% smaller than activity related to the mechanical loads (Figure 7F, I), whereas activity related to the cursor jump was 21% smaller than activity related to the mechanical loads (Figure 7G, J). Muscle activity in response to the cursor slide also gradually accumulated (Figure 7K). Cursor-slide muscle activity was 85% smaller than activity related to the mechanical loads (Figure 7L), whereas cursor-jump muscle activity was 64% smaller (Figure 7M). Collectively, these results suggest M1 and muscle display 2.9- and 6.6-times greater activities, respectively, for deviations of the hand generated by a mechanical disturbance as compared to a similar-sized visual disturbance.

## Discussion

We explored how visual and proprioceptive information related to the limb and goal are represented in M1. We found many neurons in M1 responded to sensory feedback about the limb and goal. Importantly, these different feedback sources were organized in M1 such that they largely targeted the same neurons and generated the same population-level structure.

Vision and proprioception had rapid and potent influences on M1 processing. We found a small majority of neurons responded to proprioceptive (58%) feedback consistent with previous studies (Rosén and Asanuma, 1972; Conrad et al., 1975; Lemon et al., 1976; Wong et al., 1978; Fetz et al., 1980; Lemon, 1981b; Fromm et al., 1984; Hummelsheim et al., 1988; Bauswein et al., 1991). We also found a similar percentage of neurons that responded to visual feedback of the limb (52%) and goal (55%). Both visual and mechanical disturbances required corrective responses of about 3-4cm and the corresponding activity in M1 was comparable in size to the activity that initiated the 8-10 cm reach. Proprioceptive feedback influenced M1 activity within ∼50ms of a disturbance, whereas visual feedback influenced M1 activity within ∼80ms of a disturbance. The longer delay for vision is partly due to processing time of the retina as the lateral geniculate nucleus, an area immediately downstream of the retina, responds to visual input within ∼20-30ms (Maunsell et al., 1999). In contrast, muscle spindles respond to a muscle stretch within ∼3ms (Schäfer et al., 1999) and the conduction delay to first-order thalamic nuclei are approximately 6ms (Lemon and van der Burg, 1979). Thus, sensory feedback has a potent influence on M1 processing when responding to external disturbances and it is likely that sensory errors generated during natural reaching also have a potent influence (Crevecoeur et al., 2012; Crevecoeur and Kurtzer, 2018; Takei et al., 2018).

Interestingly, the timing for proprioceptive feedback was noticeably longer than previous studies that demonstrate M1 responds within ∼20ms of a mechanical load (Evarts and Tanji, 1976; Wolpaw, 1980; Fromm et al., 1984; Boudreau and Smith, 2001; Pruszynski et al., 2014; Omrani et al., 2016). This may reflect task differences as previous studies have applied loads during posture, whereas the present study applied loads during reaching. Alternatively, the present study recorded M1 neurons using floating micro-electrode arrays and sampled only neurons on the gyrus of M1 (rostral M1). In contrast, previous studies including our own studies recorded M1 neurons using single electrodes that sampled neurons from the gyrus as well as the most caudal portion of M1 residing in the central sulcus. Previous work suggest that there are gradients along the rostral-caudal axis of M1 for anatomical and physiological features (Crammond and Kalaska, 1996, 2000; Cisek et al., 2003; Rathelot and Strick, 2009; Witham et al., 2016). Thus, faster timing may reside in neurons sampled from the caudal subdivision of M1.

Importantly, our results support the convergence hypothesis for how M1 responds to different sources of sensory feedback. First, each feedback source targeted a highly overlapping population of neurons. Second, neurons maintained their selectivity and response range for corrections across the different perturbation types. Lastly, we found a strong similarity in the population structure as principal components trained on one perturbation type captured a substantial amount of variance for the other perturbation types. The high similarity in the population structure emerged near the time when perturbation-related activity emerged suggesting that these feedback sources converged rapidly in the network. Thus, sensory feedback about the limb and goal converge onto the same circuit in M1 and give rise to similar population-level structure.

The high convergence of sensory feedback suggests that areas upstream of M1 are responsible for combining these information sources. Frontal and parietal cortices are likely involved with state estimation where proprioceptive and visual feedback are integrated into a common limb estimate (Desmurget and Grafton, 2000; Shadmehr and Krakauer, 2008; Scott, 2012; Takei et al., 2021). These areas receive proprioceptive and visual feedback with subpopulations of neurons that are responsive to both sensory modalities (Rizzolatti et al., 1981a, 1981b; Snyder et al., 1998; Bakola et al., 2010; Omrani et al., 2016; Gamberini et al., 2017). Several neurophysiological investigations have also indicated that these same areas are involved with generating a movement vector by combining limb and goal feedback (Snyder et al., 1998; Buneo et al., 2002; Pesaran et al., 2006; McGuire and Sabes, 2011; Bremner and Andersen, 2012; Piserchia et al., 2017). While this movement vector is commonly assumed to reflect a spatial representation, it may reflect a more complex neural space including information related to arm geometry (Scott et al., 1997).

Consistent with upstream state estimation is that M1 activity was largely unaffected by the transient removal of cursor feedback. Other groups also found that the motor system was insensitive to the removal of cursor feedback, but interpreted this as evidence that reaching involves a ballistic phase where feedforward motor commands transport the limb towards the goal with little influence from sensory feedback (Woodworth, 1899; Meyer et al., 1988; Suway and Schwartz, 2019). However, our perturbations show that M1 is still highly sensitive to proprioceptive and visual feedback inconsistent with this ballistic interpretation. The insensitivity to cursor visibility likely reflects that the motor system also uses internal and proprioceptive feedback to compensate for missing visual information consistent with multi-sensory state estimation (Crevecoeur et al., 2016). This compensation strategy is likely necessary as shifts in the gaze position and blinks can disrupt the visibility of the hand during motor actions. Further, we found a small increase in the distance the reach endpoint was from the goal when cursor feedback was removed suggesting only a partial compensation by these alternative feedback sources.

Although convergence upstream of M1 is likely, there are two reasons why convergence may also arise from local processing in M1. First, a difference vector by definition is a relative metric about how far the limb is from the goal and thus cannot update M1 about the current limb configuration. Information about the limb configuration is necessary for control to account for state-dependent properties of the limb (e.g. intersegmental dynamics Hollerbach and Flash, 1982; Sober and Sabes, 2003; Kurtzer et al., 2008; Pruszynski et al., 2011). Second, M1 receives direct and substantial inputs from S1 and the interpositus nuclei of the cerebellum, areas which are likely involved with state estimation and exhibit activity patterns independent of the goal (Vilis et al., 1976; Strick, 1983; Omrani et al., 2016). Local convergence of sensory feedback may arise in M1 by initial processing in layers 2/3 as these layers rapidly respond to proprioceptive and visual feedback (Lemon, 1981a; Chandrasekaran et al., 2017; Heindorf et al., 2018). Alternatively, convergence may arise from integration by the dendrites of layer 5 M1 neurons. Further studies are required to understand how sensory feedback signals are combined in frontoparietal circuits including M1.

Our results also highlight differences between corrections for mechanical and visual perturbations at the muscle and M1 levels that provide potential insight about the relative contribution of M1 in feedback processing. M1 and muscle activities were larger for the mechanical loads than sliding cursor perturbations that followed a similar kinematic trajectory. This difference in magnitude may reflect a combination of two factors. First, the motor system may only use visual feedback to update internal estimates of the kinematic variables and thus corrections are generated to counter the kinematic error only. In contrast, proprioceptive feedback may be used to update estimates of kinematic and dynamic variables including the external load and thus corrections are generated to counter both the kinematic error and the external load. Second, a sliding cursor perturbation introduces a conflict between visual and proprioceptive feedback which may have attenuated the accompanying corrective response. Multi-sensory integration theories suggest the motor system should weight proprioceptive and visual feedback to form a common limb estimate with a recent study suggesting proprioceptive feedback should be weighted more given its shorter delays compared to vision (Crevecoeur et al., 2016). In contrast, we removed cursor feedback on mechanical load trials and thus there was no conflict between vision and proprioception. Further studies are needed to probe what state variables are updated by each sensory modality and the integration rules used by the motor system.

There was also a noticeable difference in the relative magnitudes for visual and mechanical perturbations between M1 and muscle activities. Muscle activity was 6 times larger for the mechanical loads than cursor slides, whereas M1 activity was only 3 times larger for mechanical loads than cursor slides. This suggests that M1 only contributes ∼50% of the total motor output for mechanical loads with the remaining output likely generated by subcortical circuits including brainstem and spinal cord (Mewes and Cheney, 1991; Soteropoulos et al., 2012; Herter et al., 2015; Soteropoulos and Baker, 2020). However, this estimate on the cortical contribution to motor corrections has many assumptions. First, the activity we recorded in rostral M1 is assumed to be representative of descending cortical control, in general. Further studies are clearly required to verify whether the relative difference is reflective of regions such as caudal M1 in the bank of the central sulcus where proprioceptive and cutaneous responses tend to be greater (Porter and Lemon, 1993). Second, it is assumed that neural responses for visual and mechanical disturbances contribute similarly to descending signals or output-potent spaces (Kaufman et al., 2014; Stavisky et al., 2017). This assumption seems reasonable as the population-level structure was largely similar between mechanical and visual perturbations. Finally, it is likely that we underestimated the subcortical contribution to mechanical loads as the comparison between mechanical and visual perturbations assumed M1 was the only circuit involved with generating visual responses. Visual responses may also involve subcortical circuits including the superior colliculus (Alstermark et al., 1987; Day and Brown, 2001; Pruszynski et al., 2010; Corneil and Munoz, 2014; Cross et al., 2019; Kozak et al., 2019). While comparisons of the visual and mechanical responses at the muscle and neural levels provides a potentially important approach to probe cortical versus subcortical contributions to feedback corrections, further studies are clearly required to address the assumptions inherent in these estimates.

The presence of feedback processing at cortical and subcortical levels highlight that the motor system is hierarchically organized with feedback at multiple levels and transcortical feedback through M1 being the highest level for online continuous control (Porter and Lemon, 1993; Schweighofer et al., 1998; Loeb et al., 1999; Todorov et al., 2005; Liu and Todorov, 2009; Merel et al., 2019). Current theories inspired by engineering principles have adopted a serial approach focused on the transformation of information (e.g. cartesian space to joint torques; Kalaska and Crammond, 1992; Buneo et al., 2002; Todorov et al., 2005) or a modular approach where each level provides a distinct role (e.g. motor planning by motor cortex, feedback control by subcortical circuits; Kawato et al., 1987; Schweighofer et al., 1998; Loeb et al., 1999; Merel et al., 2019). Alternatively, multiple levels may contribute to generating feedback responses, but without distinct roles captured by engineering principles. From this perspective, the contribution by M1 would be to provide the extra motor commands necessary to attain a behavioural goal that is adjusted based on the expected contributions provided by lower feedback pathways. This could even include a reduction in motor output when needed to compensate for increased contributions from lower circuits (e.g. gain scaling, Pruszynski et al., 2009). Unravelling the relative contributions of different levels of the motor system during voluntary control remains an important and challenging area of study.

## Methods

The study involved two monkeys (*Macaque mulatta*, males, 17-20kgs) and was approved by the Queen’s University Research Ethics Board and Animal Care Committee. Monkeys were trained to place their upper limb in an exoskeleton robot (Kinarm, Kingston Ontario).

### Lateral reaching task

Monkeys were trained to make goal-directed reaches while countering unexpected perturbations to the limb or goal. At the beginning of a trial, the monkey placed and held their hand inside a start target (red square, length and width 1.2cm,) for 750-1500ms. Then, a goal target (white square, length and width 1.6cm; joint configuration in middle of reach: shoulder 30°, elbow 87°) appeared lateral to the starting position that indicated the spatial location of the goal and provided the cue to initiate the reach. The reach primarily involved a shoulder and elbow extension motion and for Monkeys M and A, the goal targets were placed 10cm and 8cm from the start target, respectively. Monkeys had 1400ms to reach the goal and maintain their hand inside the goal for 500ms to receive water reward. We included trials where visual feedback of the hand (white circular cursor, diameter 1.6cm) was provided for the entire trial duration and trials where visual feedback of the hand was removed 2cm into the reach and re-appeared 200ms later. On random trials, we applied one of three perturbation types, goal jumps, cursor jumps, or mechanical loads. Mechanical loads consisted of torques applied to the shoulder and elbow joints in two opposite directions, one that flexed the shoulder and extended the elbow and the other that extended the shoulder and flexed the elbow. Shoulder and elbow torques were equivalent in magnitude and were 0.28Nm and 0.24Nm for Monkeys M and A, respectively. Visual feedback of the hand was also removed for 200ms after the mechanical load was applied. Cursor jumps consisted of displacements to the cursor’s position perpendicular to the axis connecting the start and goal targets (reach axis, Figure 1A). Two cursor-jump directions were included that displaced the cursor away from or towards the body and the size of the displacement was 4cm and 3cm for Monkeys M and A, respectively. Goal jumps were identical to cursor jumps except that the goal’s position was displaced. All perturbations were applied 2cm into the reach. In a block of trials, monkeys performed 8 unperturbed reaches with visual feedback of the hand, 4 reaches with visual feedback of the hand temporally removed for 200ms and 6 perturbation trials (2 directions x 3 perturbation types). Monkeys completed 10-25 blocks in a recording session.

### Anterior reaching task

For a subset of sessions, monkeys also completed reaches to a goal located directly in front of the shoulder (anterior reach). These reaches followed the same timing parameters as the lateral reaches denoted above. Goal and cursor jumps were still in the direction that was lateral to the reach axis, which now resulted in jumps that were lateral or medial to the body. Mechanical loads were the same magnitude, however now they either flexed the shoulder and elbow joints or extended the shoulder and elbow joints. In a recording session, monkeys completed 10-15 blocks of the lateral reaches followed by 10-15 blocks of the anterior reaches or completed the anterior reaches first followed by the lateral reaches. The ordering of the blocks were counterbalanced across sessions.

### Cursor slide task

In a separate set of experiments, we probed the sensitivity of M1 activity to proprioceptive and visual stimuli when the temporal and spatial characteristics were matched. Monkeys completed the same lateral reaching task with the same mechanical load and cursor jump perturbations. However, we also included a cursor slide perturbation where the visual location of the cursor would traverse a trajectory similar to the trajectory the limb would take following a mechanical load. We estimated the trajectory by fitting the limb position on mechanical load trials to a sigmoid function (a/(exp(-(t+b)/c), where *t* is time and *a, b, c* are fit parameters) from 50ms before the load till 200ms after the load onset. The sigmoid fit parameters were estimated using trials from a previous day’s recording session.

### Estimating visual onsets

There is an approximate 20-40ms latency in the visual display between when a command is sent to jump the cursor or goal and when it appears on the screen. On a trial-by-trial basis, we estimated the visual latency by fixing two photodiodes to the screen. When the goal or cursor jumped, two white squares would also appear that were positioned on the screen coincident with the photodiode placements. Jump onsets were estimated as the average onset of the two photodiodes, or the onset detected by a single photodiode when the other photodiode signal was poor. On trials where a cursor and goal jump did not occur, the white squares still appeared at the same point in the reach so that we could align the unperturbed trials.

### Neural recordings

In each monkey, floating micro-electrode arrays (96-channel, Utah arrays) were surgically implanted into the arm region of primary motor cortex. Surgery was performed under aseptic conditions and the arm region was identified by visual landmarks. During surgery we used a dura substitute (GORE PRECLUDE Dura Substitute, W.L. Gore and Associates Inc) that was placed over the array and the dura was re-attached (GOR-TEX Suture, W.L. Gore and Associates Inc). Spike waveforms were sampled at 30 kHz by either a 128-channel neural signal processor (Blackrock Microsystems, Salt Lake City, Utah) or a Grapevine processor (Ripple Neuro, Salt Lake City, Utah). Neural recordings were collected over 5 separate recording sessions in Monkey M and 3 separate recording sessions in Monkey A.

### Muscle recordings

In Monkey M, we surgically implanted a 32-channel chronic EMG system (Link-32, Ripple Neuro, Salt Lake City, Utah). This system had 8 leads (impedance 20 kOhms) that could be inserted into the muscle with each lead having 4 separate contacts for recording muscle activity. Each lead was connected to an internal processor that was surgically implanted under the skin and located near the midline of the back at the mid-thoracic level. We implanted brachioradialis, brachialis, the lateral and long heads of the triceps, biceps (long head), pectoralis major, and anterior and posterior deltoids. During a recording session, an external transmitter was attached on the skin over the internal processor and maintained in position by a magnet in the processor. The internal processor received power from the transmitter and transmitted the EMG signals transcutaneously. The signal was transmitted to the Grapevine processor, bandpass filtered (15-375Hz) and recorded at 2 kHz. EMG recordings were collected over 3 separate recording sessions in Monkey M.

## Data Analysis

### Kinematic analysis

Kinematic signals were low-pass filtered with a 6^th^ order, zero-phase lag Butterworth filter (cut-off frequency 10Hz). The endpoint of the reach was defined as the first time point after the peak hand speed that was less than 10% of the peak hand speed. Movement time was defined as the time duration between when the monkey left the start target and first entered the goal target. We quantified the goodness of fit (R^2^) of the cursor slide trajectories (P_curs_) to the mechanical limb (P_mech_) by taking the limb position from 0-200ms after the perturbation onset and subtracting off the mean limb positions for each. We then calculated the R^2^ = 1-||P_curs_-P_mech_||^2^/||P_mech_||^2^ where ‘|| ||’ is the Frobenius norm.

### EMG recordings

Muscle activity was down sampled to 1kHz. For a given lead, we computed the differential signals between the two most proximal contacts and the two most distal contacts resulting in two differential signals from each recorded muscle. The differential signals were rectified and smoothed with a Butterworth low-pass filter with zero-phase lag at a cut-off frequency of 50Hz. Muscle activity was aligned to perturbation onset or the equivalent onset on unperturbed trials and trial averaged. For muscle activity related to mechanical perturbations, we subtracted the activity on unperturbed reaches without visual feedback from the activity on mechanical perturbation reaches. For activity related to the visual perturbations, we employed the same method except using activity on unperturbed reaches with visual feedback. The muscle’s preferred perturbation direction was determined for each perturbation type by calculating the activity with the largest perturbation response within the first 300ms of the perturbation onset. Activity was normalized by the mean activity in the first 300ms after the perturbation onset for each muscle signal.

### Pre-processing neural recordings

Spike timestamps were convolved with a kernel approximating a post-synaptic potential (1ms rise time, 20ms fall time; Thompson et al., 1996) to estimate the instantaneous activities. Activities were aligned to perturbation onset following the same procedure as for muscle activities

### ANOVA analysis

For each neuron/muscle we applied a 3-way ANOVA with time epoch (levels: baseline epoch -100-0ms, perturbation epoch 0-300ms), perturbation direction (two levels) and perturbation type (levels: mechanical loads, goal jumps and cursor jumps) as factors. Neurons/muscles were classified as “perturbation responsive” if there was a significant main effect for time, or any interaction effects with time (p<0.05, Bonferroni correction factor=4). Neurons/muscles classified as significant were then subjected to separate two-way ANOVAs for each perturbation type with time and direction as factors. Neurons/muscles were classified as responsive for a given perturbation type if a significant main effect or interaction effect was found (p<0.05, Bonferroni correction factor=2).

### Response range

The response range for a neuron was calculated for each perturbation type separately by taking the activity related to the correction towards the body and subtracting the activity related to the correction away from the body. The resulting activity was then averaged over the perturbation epoch.

### Total least-square (TLS) regression

TLS regression was used to find a linear relationship between the response ranges from two perturbation types (Figure 5). Ordinary least square (OLS) regression has been used in previous studies (Crammond and Kalaska, 2000), however, this method assumes one set of response ranges is the independent variable (i.e. no sampling noise; denote as x) and thus only tries to find a line that minimize the error between the dependent variable (y) and the line (minimize 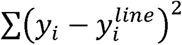). In contrast, TLS regression does not assume any variables are independent and finds a line of best fit that minimizes the total error between each data point and the line (minimize 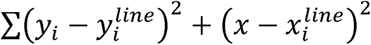). TLS was performed by first subtracting the means for each response range 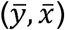 followed by singular value decomposition to find the slope (m). The left singular vector with the largest singular value was retained and the slope of the line of best fit was given as the ratio between the coefficient for the data on the y-axis over the coefficient for the data on the x-axis. The equation of the line of best fit is then *y*^*line*^ = *m* · *x*^*line*^ + *b* where 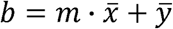. The significance of the slope was determined by shuffling the perturbations labels and re-calculating the slope. This was repeated 1000 times and a probability value was calculated as the number of shuffled samples with slope smaller than the actual slope.

### Onsets

The onset of perturbation-related activity was estimated by calculating the mean and standard deviation of the perturbation-related activity during the baseline period (100ms before perturbation onset). The onset was then defined as the first time-point to exceed the baseline mean by three standard deviations (positive or negative) for 20 consecutive time points. This method was used to calculate the onset for individual neurons, the neural population activity and the muscle population activity. For individual neurons, the onset was only calculated once per neuron in the perturbation direction that elicited the largest absolute response from the unperturbed trials during the perturbation epoch.

### Average population activity

An average population response was calculated to estimate the total change in the network in response to the perturbations. We determined each neuron’s preferred corrective movement by averaging its activity over the perturbation epoch. The corrective movement with the absolute largest change in activity from the unperturbed activity was then defined as the preferred corrective movement. If the change in activity was negative for a neuron in its preferred corrective movement, we multiplied its time series by negative one. This reduced the cancelling out of activity when averaging across the population of neurons.

### Principle components analysis

Principal components analysis (PCA) was used to identify the low-dimensional subspace for the perturbation-related activity. For each perturbation type, we averaged each neuron’s perturbation-related activity in non-overlapping 10ms windows to yield 30 time points for each perturbation direction. The activity of each neuron was soft normalized by its range (+5 sp/s) by finding its maximum and minimum activities during the perturbation epoch over all perturbation types (mechanical loads, goal jumps, cursor jumps). Note, the same normalization constant was applied to each perturbation type. We then constructed separate matrices for each perturbation type that were of size NxDT, where N is the number of neurons, D is the number of perturbation directions (2) and T is the number of time points (30). The mean activities in each row was then subtracted. Singular value decomposition was used to identify the principle components of the matrix, and the top-10 principle components were kept.

We used a cross-validated approach to draw a more accurate comparison between the amount of variance captured between perturbation types. For a given perturbation type, we randomly assigned trials into equally sized groups and the same processing steps were applied as above. One group was used to calculate the principle components (Trained) while the left-out group was used to calculate the amount of variance captured by those principle components. These principle components were also used to calculate the amount of variance accounted for by the other two perturbation types after randomly down-sampling trials to match the left-out group. This procedure was repeated 1000 times for each perturbation type.

### Overlap index

We quantified the overlap between the subspaces by calculating the overlap index from Rouse and Schieber, (2018)

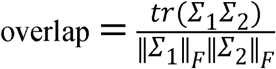

Where Σ_l_ and Σ_2_ are the covariance matrices for perturbation types 1 and 2, *tr* is the trace operator, and ‖ · ‖_*F*_ is the Frobenius norm operator. Activity was pre-processed the same way as for the PCA analysis. The overlap index was computed between each pair of perturbation types.

The overlap index can range from 0, indicating no overlap between subspaces, and 1 indicating perfect overlap between subspaces. Confidence intervals were generated by randomly selecting half of the trials for each perturbation condition and calculating the subsequent overlap. This was repeated 1000 times for each comparison between perturbation types.

We generated two null distributions for comparison. One distribution estimated the overlap between two independent samples from the same perturbation type (within-perturbation distribution). For a perturbation type, we split trials into two, equally sized groups and then calculated the overlap between these two groups following the same procedure as above. This was repeated 1000 times for each perturbation type and overlap values were pooled. The second distribution compared how overlapping two samples were when the neuron labels were shuffled. For a perturbation type, we again split trials into two, equally sized groups. The neuron labels were then randomly shuffled in one group and the overlap was then calculated between the two groups. This was repeated 1000 times for each perturbation type and overlap values were pooled.

## Acknowledgements

We thank Kim Moore, Simone Appaqaq, Ethan Heming, and Helen Bretzke for their laboratory and technical assistance and the LIMB lab for helpful discussions. This work was supported by grants from the Canadian Institute of Health Research. KPC was supported by an OGS scholarship. SHS was supported by a GSK chair in Neuroscience.

## Figures Legends

**Supplementary Figure 1.**
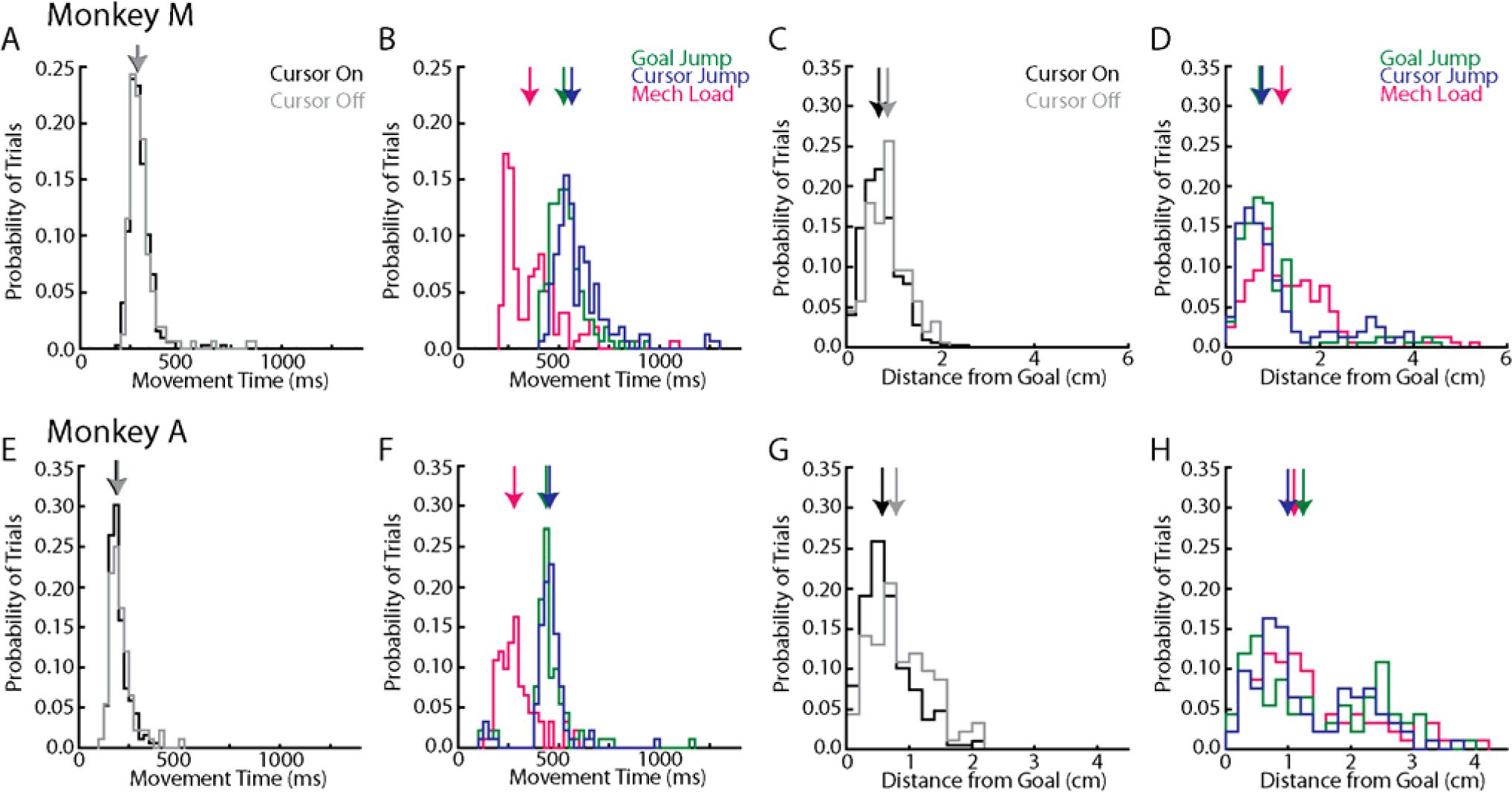
Movement times and endpoint distance from goal across monkeys. A) Movement times for Monkey M for cursor-on and cursor-off unperturbed reaches. Movement time was defined as the time between when the hand left the start target and when the hand first contacted the goal target. Trials have been pooled across all recording sessions. Arrows denote the median of the distributions. Distributions for cursor-on and cursor-off trials were not significantly different (two-sample t-test: t(471)=1.6, p=0.12). B) Same as A) for perturbation trials. C) Same as A) except for the distance the reach endpoint was from the goal. Distributions for cursor-on and cursor-off trials were significantly different (t(471)=3.6, p<0.001). D) Same as C) for perturbation trials. E-H) Same as A-D) for Monkey A. E) Distributions for cursor-on and cursor-off trials were not significantly different (t(279)=1.9, p=0.06). G) Distributions for cursor-on and cursor-off trials were significantly different (t(279)=4.0, p<0.001). Note, Monkey M had longer movement times than Monkey A due in part to Monkey M completing a 10cm reach and Monkey A completing an 8cm reach.

**Supplementary Figure 2.**
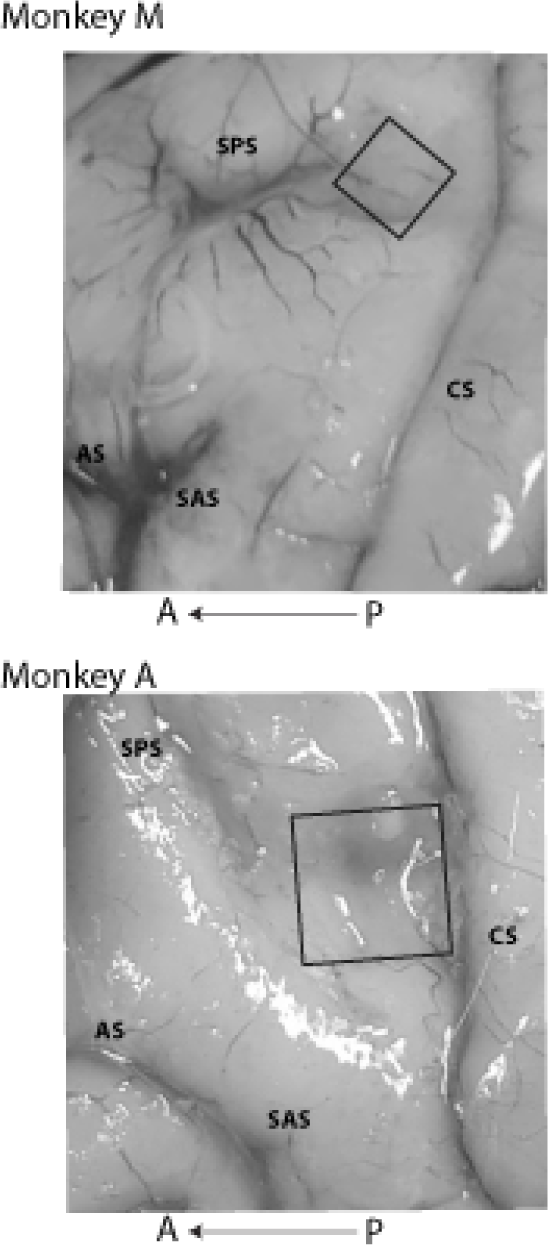
Placement of arrays in M1. Both monkeys had floating micro-electrode arrays implanted in the arm regions of M1. Approximate location of array is indicated by the black square. Acronyms: CS central sulcus, SPS superior precentral sulcus, AS arcuate sulcus, SAS spur of arcuate sulcus, A anterior, P posterior.

**Supplementary Figure 3.**
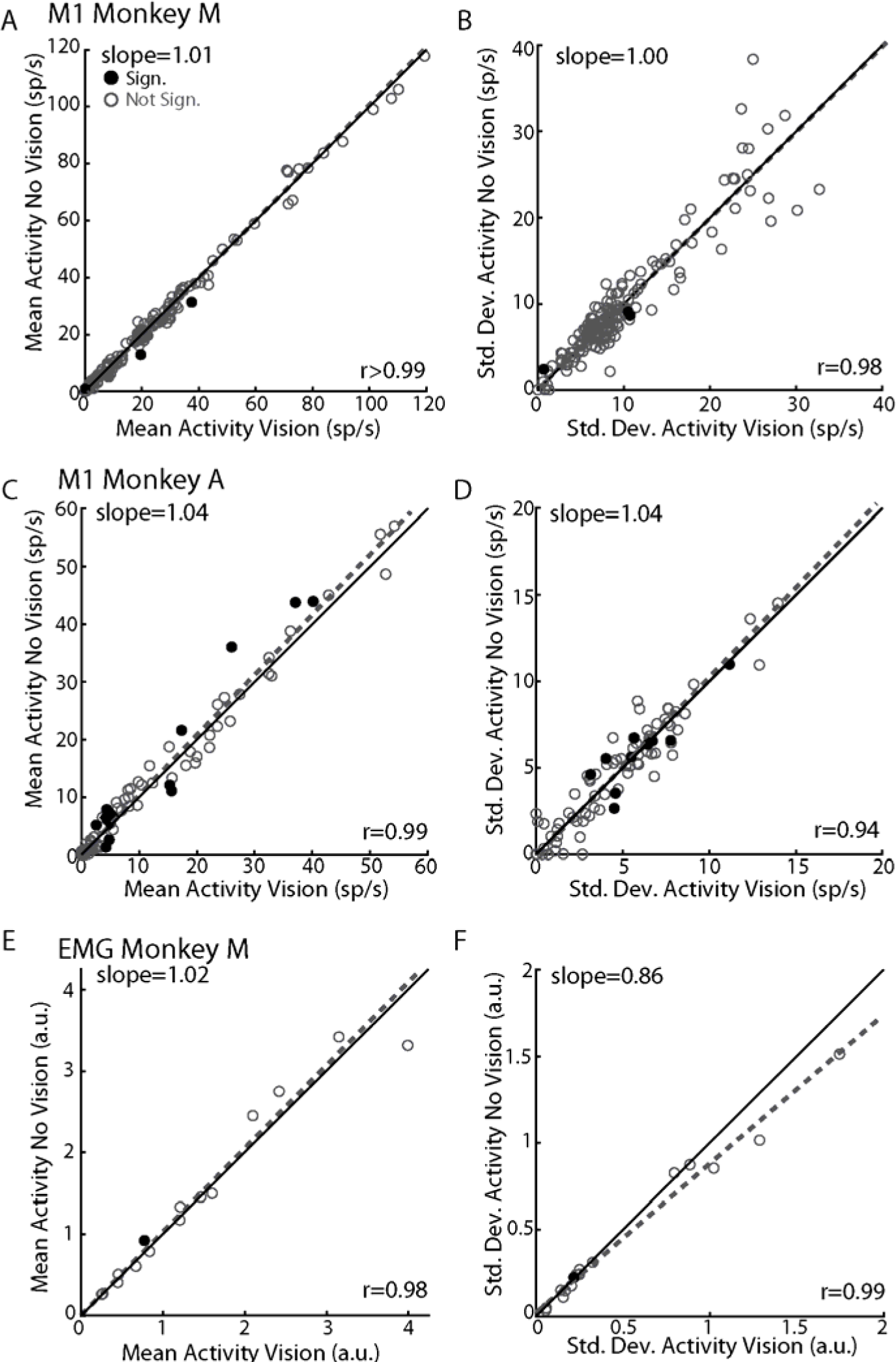
M1 activity is largely unaffected by removing cursor feedback. A) For Monkey M, comparison of the mean activities during unperturbed reaches for cursor-on (abscissa) and cursor-off (ordinate) trials. Activity was averaged from 100-250ms after the cursor feedback was removed. Each circle denotes one neuron. Dashed line reflects the line of best fit identified using total least squares regression (slope indicated in top left corner). B) Same as A) except for the standard deviation across trials. C-D) Same as A-B) except for Monkey A. E-F) Same as A-B) except for EMG from Monkey M.

**Supplementary Figure 4.**
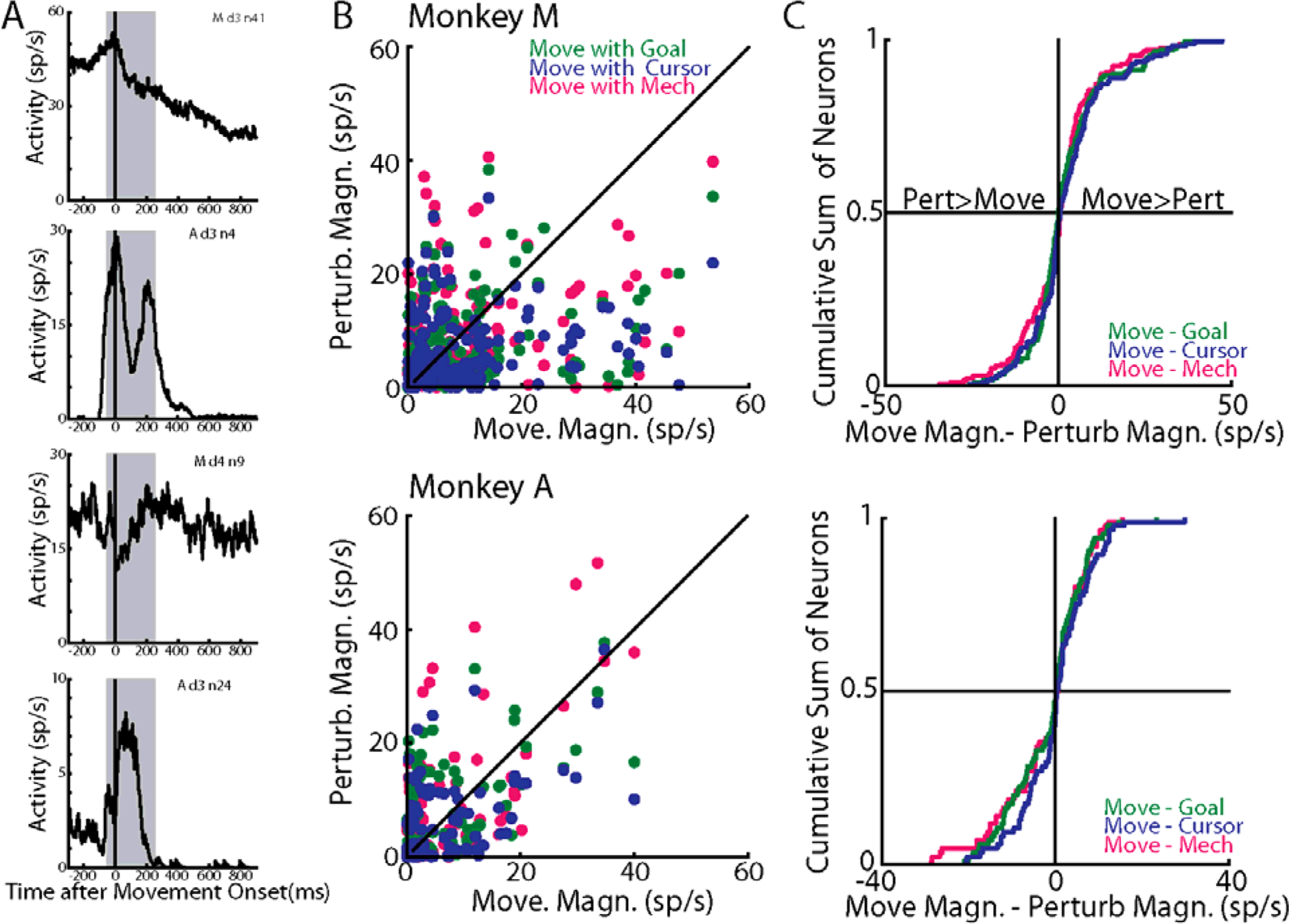
Perturbation-related activity is comparable to activity during baseline reaching. A) Activities of the same four example neurons in Figure 2 during unperturbed reaches aligned to movement onset (5% max hand speed). Shaded area denotes the movement epoch (−50-250ms). B) Scatter comparing the absolute magnitude of movement-related activity with the magnitude of the perturbation-related activity. C) Cumulative sums of the difference in the magnitudes of the movement-related and perturbation-related activities across cells.

**Supplementary Figure 5.**
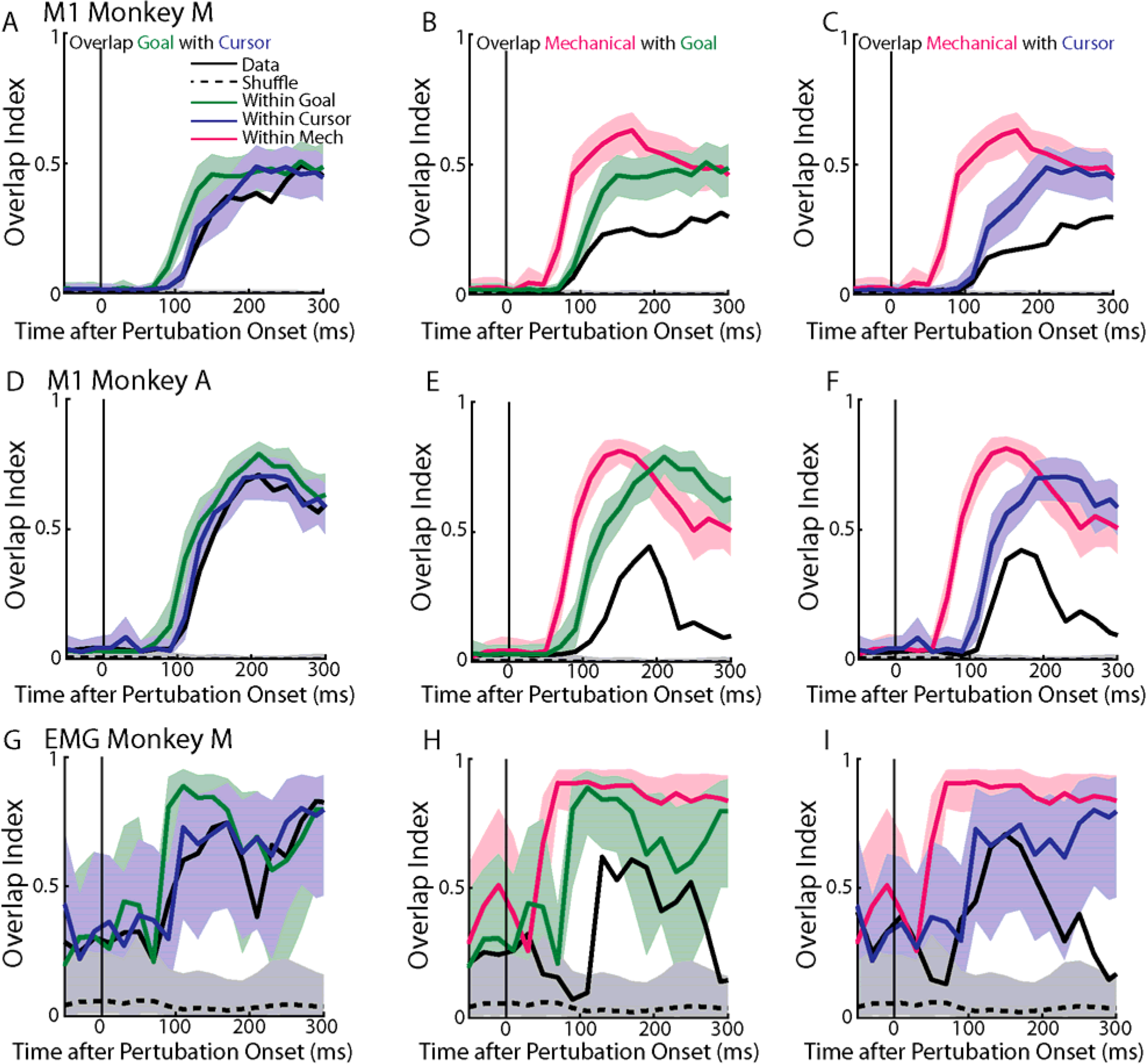
Overlap time course. A) Time series of the overlap index between goal and cursor jumps (black solid line) for Monkey M. Activity was binned every 20ms. The time series was also repeated for the shuffle distribution (black dashed line) and the within-perturbation distributions for the goal-related (green line) and cursor-related (blue line) activities. B) Same as A) except comparing mechanical loads with goal jumps. C) Same as A) except comparing mechanical loads with cursor jumps. D-F) Same as A-C) except for Monkey A. G-I) Same as A-C) for EMG signals. Prior to overlap calculation, EMG signals were filtered with a low-pass 3^rd^ order Butterworth filter (cut-off 50Hz). Note, the substantial overlap before perturbation onset is in part due to the small subspace spanned by EMG signals

**Supplementary Figure 6.**
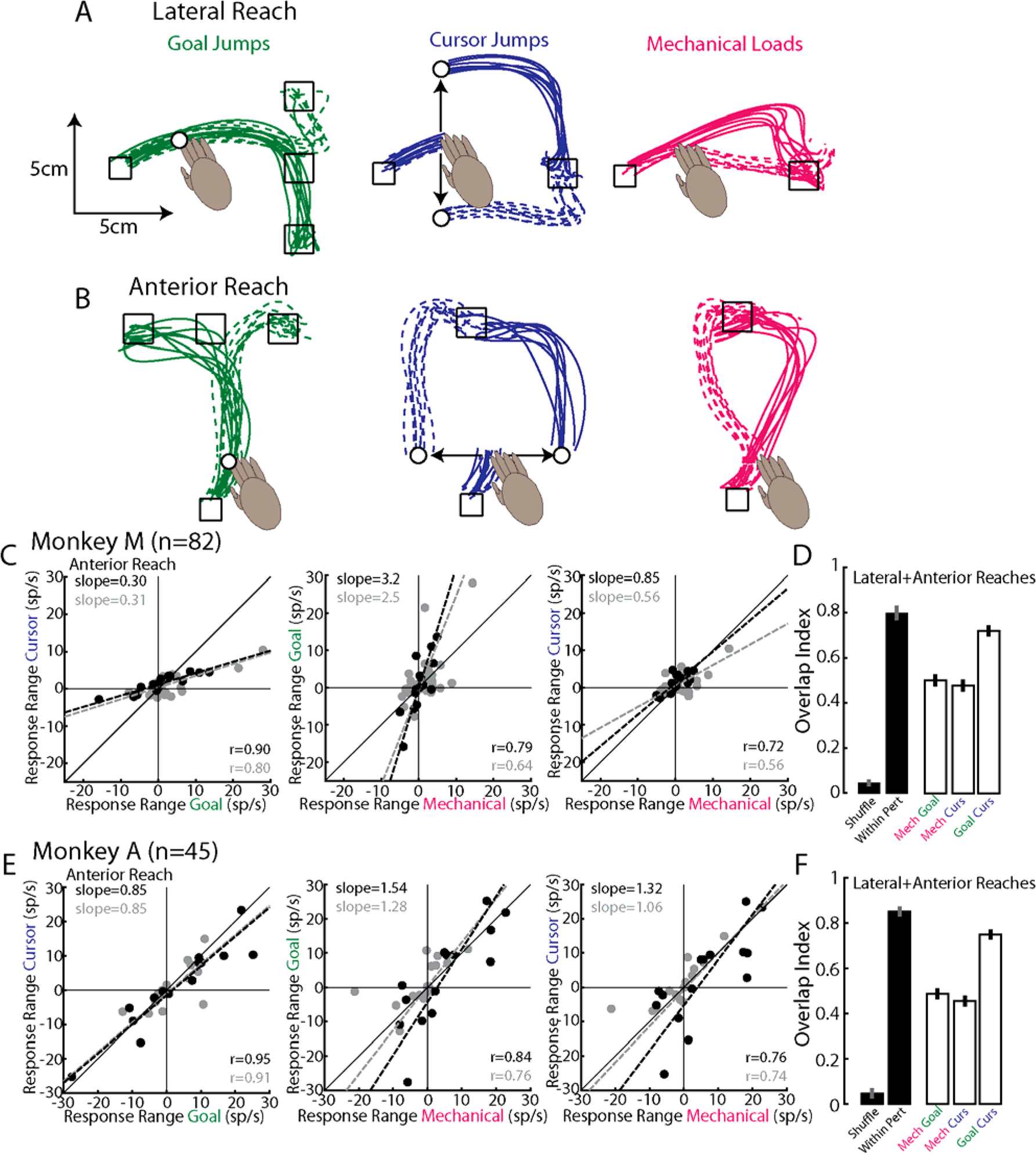
Overlap across perturbation types with increased perturbation directions. A) Monkey M’s lateral reaches following goal jumps (left), cursor jumps (middle) and mechanical loads (right). Same as Figure 1B-D. B) Same as A) except now for Monkey’s M anterior reaches. C) Response ranges comparing perturbation types for the anterior reaches. Data presented the same as in Figure 5. ‘n’ denotes the number of recorded neurons. D) Overlap index presented the same as Figure 6. E-F) Same as C-D) for Monkey A.

